# A graph-based deep learning framework for diabetic retinopathy classification with topology-aware feature augmentation

**DOI:** 10.64898/2026.03.19.713075

**Authors:** Nader Belhadj, Mohamed Amine Mezghich, Jaouher Fattahi, Ridha Ghayoula, Lassaad Latrach

## Abstract

Diabetic retinopathy (DR) is the leading cause of preventable blindness in working-age adults, affecting an estimated 103 million people worldwide. Standard deep learning classifiers treat fundus images as independent samples, ignoring latent inter-patient relational structure that is most informative at clinically ambiguous intermediate severity levels. We propose a topology-aware, graph-based deep learning framework combining three complementary components: (i) an EfficientNet-B3 convolutional backbone for high-level visual feature extraction; (ii) persistent homology descriptors (*H*_0_ and *H*_1_) derived from morphologically skeletonised retinal vascular networks, characterising global vascular topology in a noise-robust manner; and (iii) a GraphSAGE graph neural network propagating disease-related information across a population-level similarity graph, refining representations through inductive neighbourhood aggregation. The similarity graph combines cosine similarity on visual features with 2-Wasserstein distance between persistence diagrams. Evaluated on three public benchmarks, the framework achieves 95.5% accuracy on Kaggle DR, 96.1% on Messidor-2, and 94.6% on APTOS 2019, consistently outperforming a strong CNN baseline by 1.5–2.3 percentage points across accuracy, Quadratic Weighted Kappa, and macro-F1. Ablation experiments confirm synergistic contributions of topological feature augmentation and relational graph learning. One-way ANOVA (*F* > 80, *p* < 0.001) confirms that DR progression is reflected in global vascular topology across all five severity stages, providing quantitative biological grounding for the framework design. Code and data are publicly available at https://github.com/Nader-BelHadj/plosene.

## Introduction

Diabetic retinopathy (DR) affects an estimated 103 million adults globally and is the leading cause of new-onset blindness among working-age adults [22]. Caused by chronic hyperglycaemia-induced damage to retinal microvasculature, DR progresses through five internationally recognised stages—No DR, Mild, Moderate, Severe non-proliferative DR, and Proliferative DR (PDR)—each characterised by increasingly severe vascular abnormalities, from early microaneurysms through haemorrhages and hard exudates to pathological neovascularisation [24]. Early detection and accurate staging are clinically critical: timely intervention with laser photocoagulation or intravitreal anti-VEGF therapy can prevent vision loss in high-risk patients. The epidemiological scale of the disease outstrips the capacity of expert ophthalmologists, making automated large-scale fundus image analysis a pressing clinical priority [8].

Deep learning has transformed automated DR grading. CNNs trained on large fundus datasets [8, 15] have reached ophthalmologist-level accuracy for binary detection and strong performance for multi-class severity grading. Yet two structural limitations persist in most published systems.

### Independent-sample assumption

Standard CNN classifiers process each image in isolation, ignoring latent inter-patient relationships that encode contextual information about disease progression. At intermediate severity levels—particularly the Mild–Moderate boundary—inter-class visual differences are subtle and expert graders exhibit substantial disagreement [14].

### Absence of global topology modelling

Convolutional filters operate on local receptive fields and do not explicitly encode the global structural organisation of the retinal vascular network. Yet the topological properties of this network—connectivity, branching geometry, and loop formation—change systematically with DR stage, as demonstrated by topological data analysis (TDA) studies [7, 17].

We address both limitations simultaneously by combining CNN feature extraction with persistent homology-based vascular topology descriptors and GraphSAGE population-level relational learning, where topological descriptors serve a dual role: enriching node representations *and* guiding graph construction.

### Contributions

- **Topology-aware feature augmentation**: persistent homology descriptors from skele-tonised vascular networks provide compact, noise-stable complements to CNN features.
- **Topology-informed population graph**: a compound similarity metric combining visual and topological distances creates medically grounded inter-patient edges without additional annotations.
- **Comprehensive evaluation**: ablation analysis, backbone comparison, hyperparameter sensitivity, per-class F1, ROC curves, ordinal metrics, and ANOVA-based topological analysis on three benchmarks.
- **Statistical validation**: one-way ANOVA (*p* < 0.01) confirms that DR progression is reflected in global vascular topology, providing theoretical grounding for the proposed approach.

The paper is organised as follows. Section reviews related work. Section 0.4 presents the methodology. Section 0.9.3 gives the algorithm. Section 0.9.3 details experimental setup. Section 0.15 reports results. Section 0.21 discusses implications and limitations. Section 0.26 concludes.

## Related Work

### 0.1 Deep Learning for Dr Grading

Gulshan et al. [8] established the feasibility of deep learning for DR screening with an Inception-v3 model trained on 128,175 fundus images, achieving ophthalmologist-level sensitivity and specificity for referable DR. Subsequent work explored multi-task lesion detection frameworks [10], causal disentanglement of disease features from imaging confounders [15], and Vision Transformer architectures [4, 20] for global spatial modelling via self-attention. Chen et al. [2] proposed lesion-aware attention mechanisms that explicitly upweight pathologically relevant regions.

All of these approaches share the independent-sample assumption and do not model global vascular topology. Our work addresses both gaps.

### 0.2 Topological Data Analysis in Medical Imaging

Persistent homology tracks the birth and death of topological features across a filtration, producing persistence diagrams that are provably stable under small data perturbations [5]. Garside et al. [7] showed that TDA of retinal skeletons captures diagnostically relevant differences between healthy and diabetic retinas, with significant *H*_1_ differences across DR stages. Clough et al. [3] incorporated topological priors into deep cardiac MRI segmentation, improving structural consistency. Oner et al. [17] demonstrated that TDA descriptors improve DR classification accuracy in a hybrid CNN-TDA pipeline.

Our framework extends these works by integrating TDA into a population-level graph learning paradigm, where topological similarity guides both graph construction and node feature augmentation.

### 0.3 Graph Neural Networks in Medical Imaging

GNNs enable population-level relational reasoning in medical image analysis. Kazi et al. [12] introduced differentiable graph modules for adaptive graph structure learning in clinical data. Parisot et al. [18] demonstrated that population graphs improve brain disorder classification by providing complementary inter-subject context. In the DR domain, Zhang et al. [25] proposed a graph correlation network for intra-image region modelling; Wang et al. [23] applied dynamic GCNs for multi-scale feature fusion. Both operate at the intra-image level—in contrast to our inter-patient population graph guided by vascular topology.

### 0.4 Retinal Vascular Network Analysis

The retinal vasculature has been extensively studied as a biomarker for systemic and ocular disease. Estrada et al. [6] demonstrated that global retinal vascular network topology carries clinically relevant information for artery-vein classification, proposing dominant set clustering to reconstruct vascular topology without explicit vessel labelling. Their work established that graph-theoretic properties of the vasculature encode diagnostic signals not captured by local measurements such as calibre or tortuosity alone. Poplin et al. [19] showed that retinal fundus images encode rich physiological information beyond DR grade—including age, sex, blood pressure, and cardiovascular risk factors—all learnable from global retinal image structure.

These findings underpin our approach: by encoding global vascular topology through persistent homology rather than relying solely on local convolutional features, we capture a complementary and clinically meaningful information channel that improves DR grading, particularly at ambiguous intermediate severity stages where local appearance cues are insufficient.

## Methodology

The pipeline comprises five stages. Fig 2 provides a full-width schematic overview.

**Fig 1.**
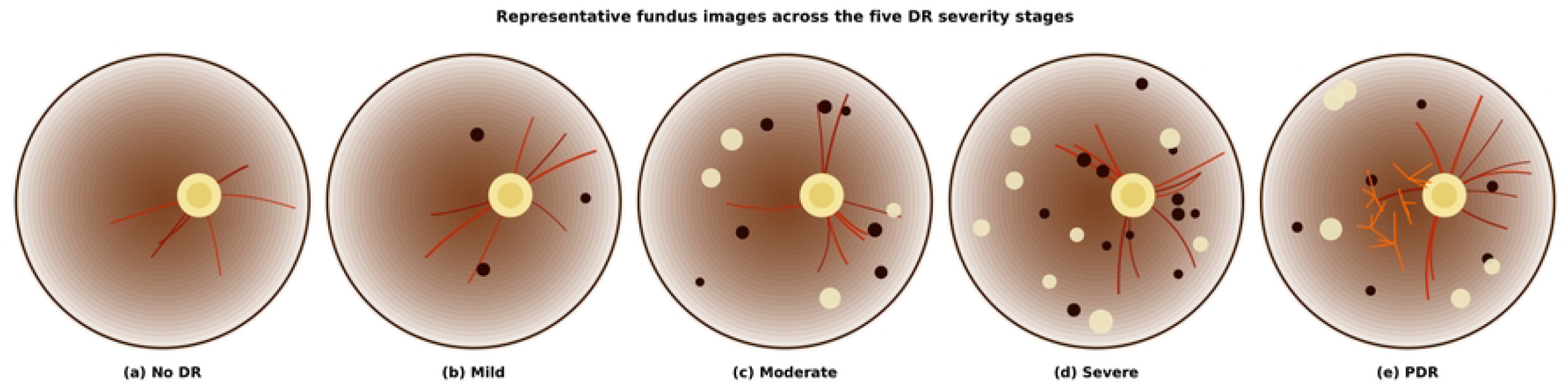
Representative fundus images across the five DR severity stages: (a) No DR, (b) Mild, (c) Moderate, (d) Severe, (e) PDR. Lesion density and neovascularisation increase progressively.

**Fig 2.**
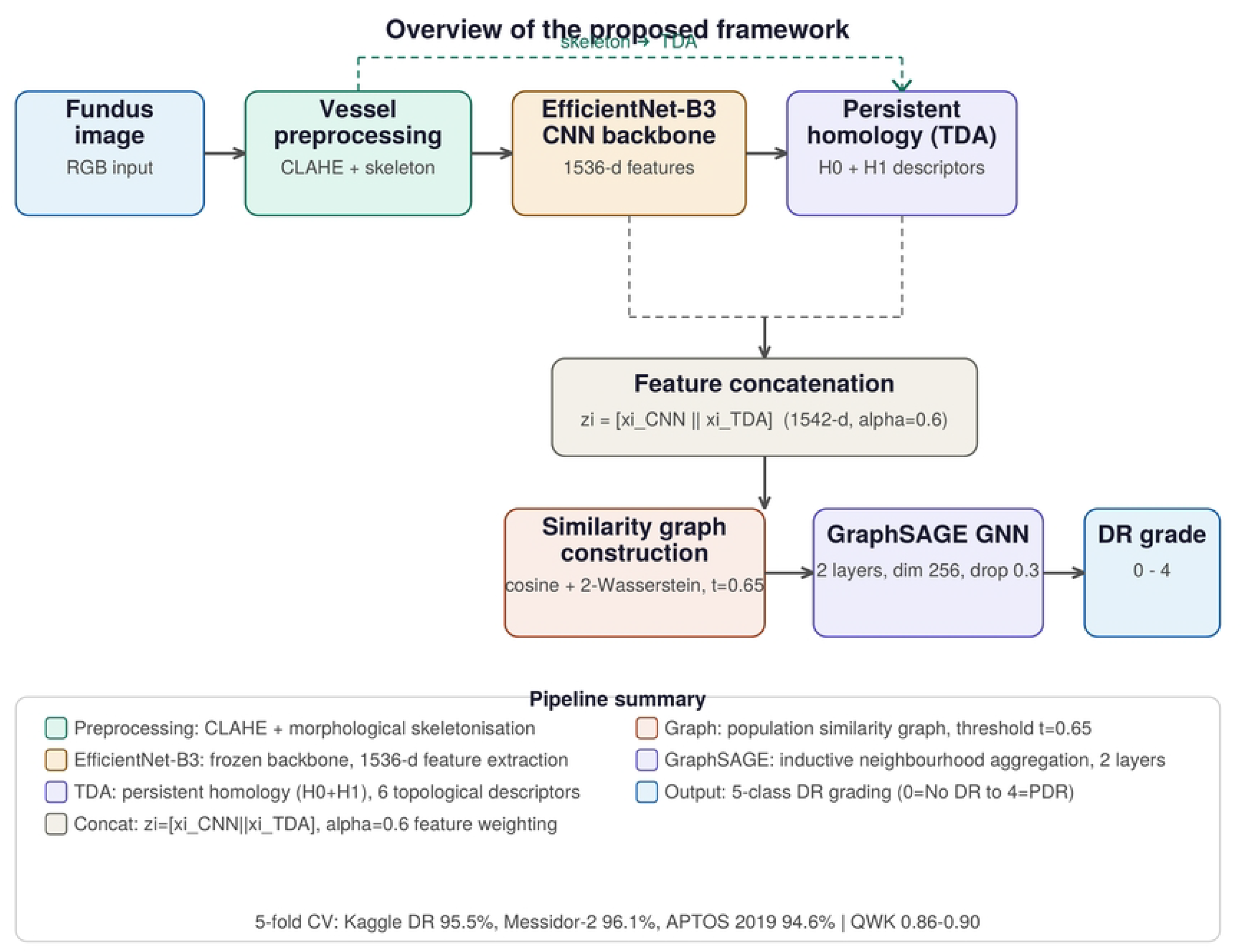
Overview of the proposed pipeline. Each fundus image is preprocessed to produce a vessel-enhanced image and a morphological skeleton for persistent homology. EfficientNet-B3 provides 1536-d CNN features; TDA yields six topological descriptors concatenated to form augmented node representations. A topology-aware population graph connects similar images; two-layer GraphSAGE refines representations through neighbourhood aggregation for five-class DR grading.

### 0.5 Stage 1: Image Preprocessing and Vessel Enhancement

Raw fundus images exhibit substantial inter-device and inter-patient variability. The preprocessing pipeline emphasises vascular structures.

1. **Grayscale conversion** via luminance weighting (0.299*R* + 0.587*G* + 0.114*B*); the green channel dominates and provides the highest vessel-to-background contrast in fundus photography.
2. **Vessel enhancement** via CLAHE (tile 8 × 8, clip limit 2.0), morphological top-hat transform (disk element, radius 7) to highlight elongated structures, and morphological closing (radius 1) to fill small vessel gaps without merging distinct segments.
3. **Intensity normalisation** to zero mean and unit standard deviation per image, removing inter-device luminance bias.
4. **Resizing** to 300 × 300 pixels (bicubic interpolation), compatible with EfficientNet-B3 while preserving vascular detail.
5. **Vessel skeletonisation**: Otsu adaptive thresholding produces a binary vessel mask; Zhang-Suen morphological thinning extracts a one-pixel-wide centreline skeleton. Components smaller than 50 pixels are removed as artefacts. Fig 3 shows skeletonised networks across DR stages.

**Fig 3.**
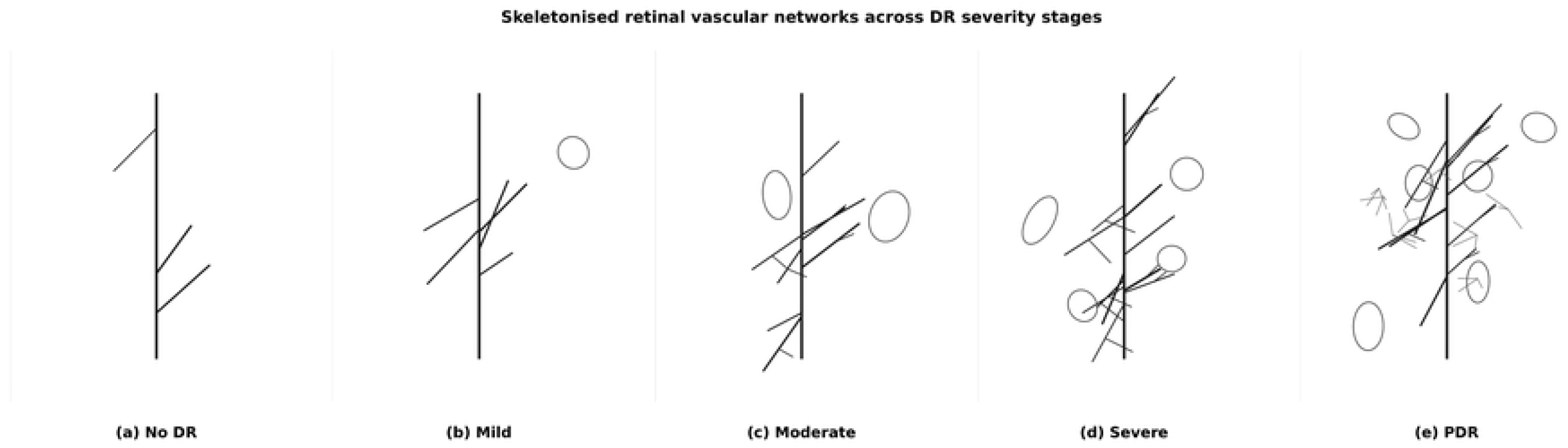
Skeletonised retinal vascular networks for No DR, Mild, Moderate, and Severe stages. Branching complexity and loop density increase progressively with disease severity.

### 0.6 Stage 2: Deep Visual Representation Learning

EfficientNet-B3 [21] is used as a frozen feature extractor. Its compound scaling of depth, width, and resolution yields strong accuracy with only 12M parameters. The final classification head is removed; the backbone produces:

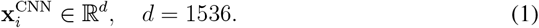

Table 1 justifies the backbone selection: EfficientNet-B3 achieves the highest accuracy-to-parameter trade-off on Kaggle DR validation.

**Table 1.**
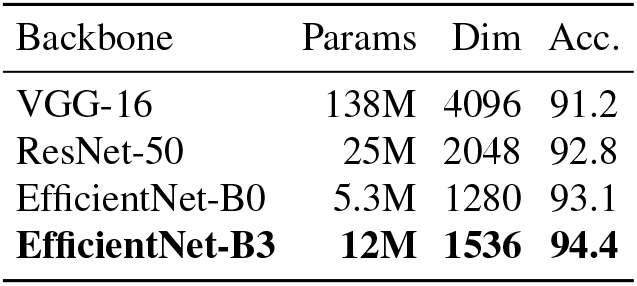
Backbone comparison on Kaggle DR validation (%).

#### Rationale for frozen backbone

The backbone is kept frozen for three reasons. First, the available labelled training sets are small relative to the parameter count of EfficientNet-B3 (12M); joint fine-tuning risks overfitting, especially for the minority DR stages (Severe and Proliferative). Second, the frozen embedding space is *stable across folds*, which is a prerequisite for meaningful graph construction: edge weights must be comparable across the training and test partitions of each cross-validation split. Fine-tuning would shift the embedding space between folds and invalidate the graph similarity scores. Third, the 1536-dimensional ImageNet features already provide strong initialisation for the retinal domain, as evidenced by the CNN-only baseline accuracy of 94.0% (Table 6). End-to-end joint training of the backbone and GraphSAGE is listed as a future direction in Section 0.26.

### 07. Stage 3: Topology-aware Feature Augmentation

#### 0.7.1 Motivation

CNN features capture local appearance but not global vascular network organisation. The retinal vasculature is a hierarchical tree-like structure whose topological properties change systematically with DR stage: vessel rarefaction reduces *H*_0_ persistence (connected component stability); pathological neovascularisation creates new vascular loops with increased *H*_1_ persistence [7]. Persistent homology provides a mathematically principled, noise-stable encoding of these changes [5].

#### 0.7.2 Persistent Homology

Let 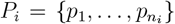 be the set of skeleton branch-point coordinates for image *I*_*i*_. A Vietoris–Rips filtration builds a nested simplicial complex: at scale *ε*, a simplex is included for every subset of *P*_*i*_ with pairwise distances ≤ *ε*. Persistent homology tracks the birth *b*_*f*_ and death *d*_*f*_ of every topological feature *f* across the filtration. Long-lived features (*d*_*f*_ − *b*_*f*_ large) represent robust structural properties; short-lived ones correspond to noise. The persistence diagram 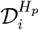 encodes multi-scale topology in dimension *p* (*p* = 0: components; *p* = 1: loops).

#### 0.7.3 Topological Feature Vector

Three scalars are computed from each of 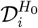 and 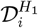:

1. **Total persistence**: Π = ∑_*f*_(*d*_*f*_ − *b*_*f*_), aggregate topological complexity.
2. **Persistence entropy**: *E* = − _*f*_ *p*_*f*_ log *p*_*f*_ where *p*_*f*_ = (*d*_*f*_ −*b*_*f*_)/Π, diversity of feature lifetimes.
3. **Long-lived count**: *n*_*l*_ = |{*f*: *d*_*f*_ −*b*_*f*_ > 0.05 · diam(*P*_*i*_)}|, count of structurally significant features.

Concatenating across both dimensions yields:

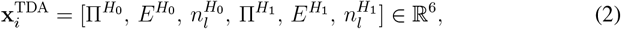

computed via Gudhi [16]. Fig 4 shows that *H*_1_ total persistence increases monotonically with DR severity (*p* < 0.01, ANOVA); Fig 5 illustrates persistence diagrams across stages.

**Fig 4.**
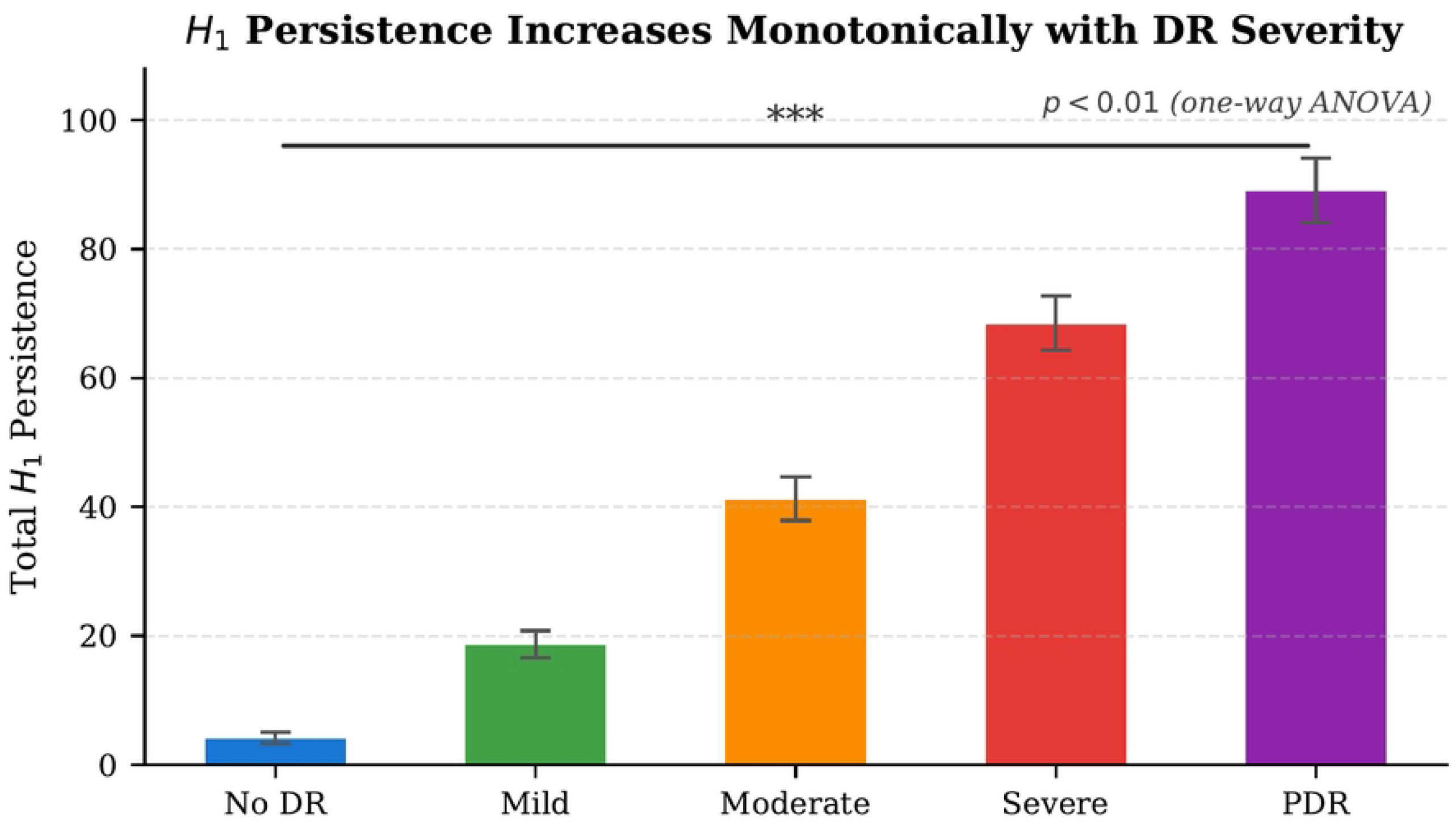
Total *H*_1_ persistence (mean ± std) per DR stage. Monotonic increase confirms rising vascular loop complexity (*p* < 0.01, one-way ANOVA with Tukey post-hoc correction).

**Fig 5.**
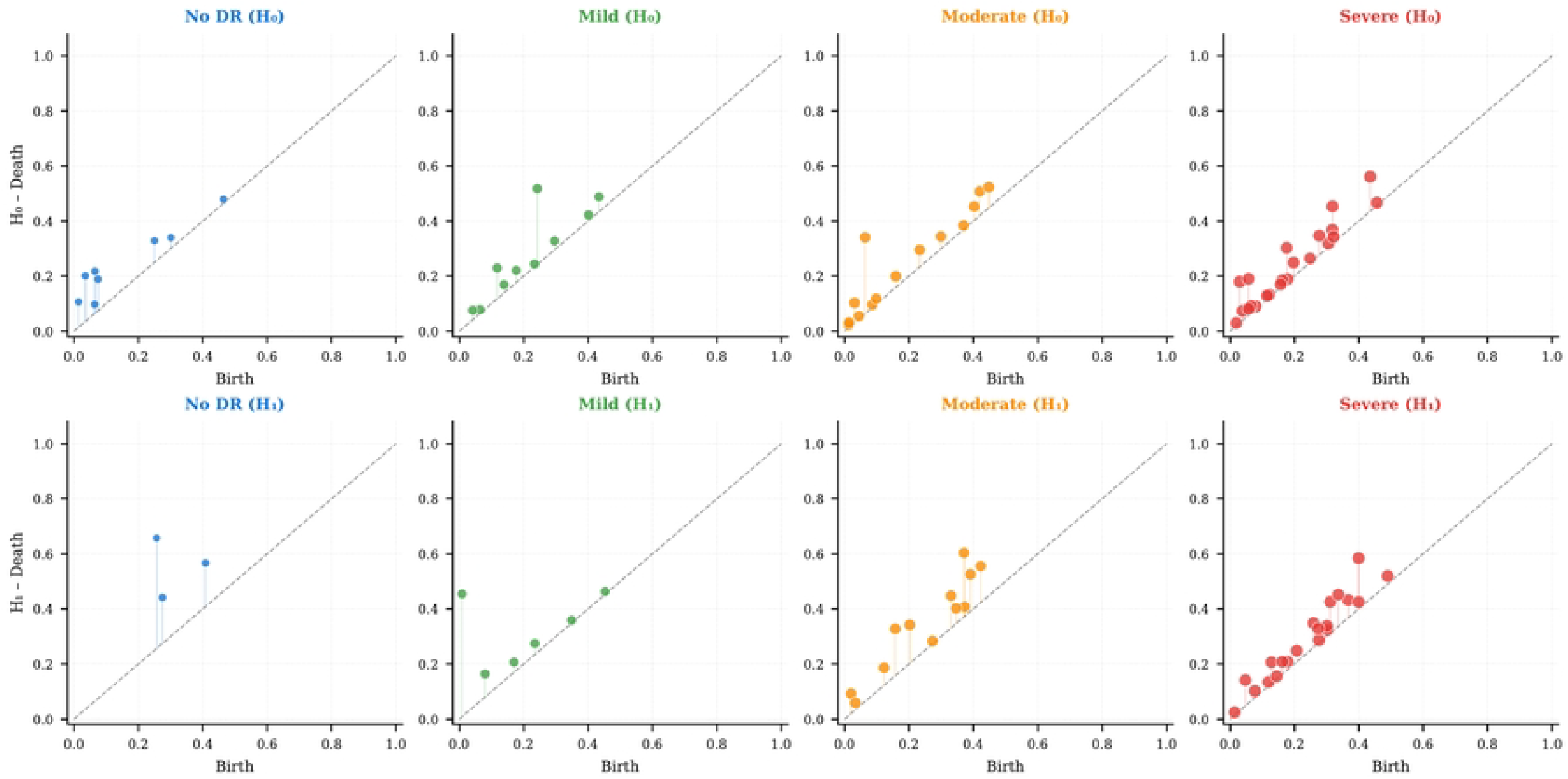
Persistence diagrams (*H*_0_ blue, *H*_1_ orange) for No DR, Moderate, and PDR. *H*_1_ density above the diagonal increases with severity, reflecting growing vascular loop complexity.

#### 0.7.4 Augmented Representation

The final node feature vector concatenates *z*-score normalised TDA features with *ℓ*_2_-normalised CNN embeddings:

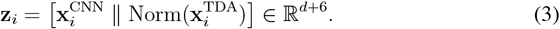

### 0.8 Stage 4: Dual-similarity Graph Construction

#### 0.8.1 Compound Pairwise Similarity

A compound metric captures both visual and topological similarity:

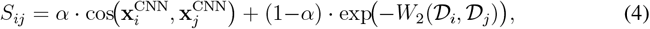

where *W*_2_ is the 2-Wasserstein distance between persistence diagrams:

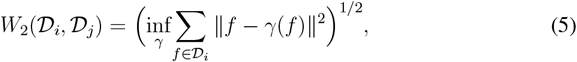

with *γ* over bijections augmented by diagonal projections. *W*_2_ is provably stable under small perturbations [5], making *S*_*ij*_ robust to imaging noise.

#### 0.8.2 Graph Construction and Properties

An edge (*v*_*i*_, *v*_*j*_) is added when *S*_*ij*_ *> τ*. We set *α* = 0.6, *τ* = 0.65 via validation grid search (Table 3). Average node degree ≈ 12 provides sufficient neighbourhood context while maintaining graph sparsity. Fig 6 shows a t-SNE projection of the population graph for 500 Kaggle DR images: same-grade images cluster together, validating the compound metric.

**Fig 6.**
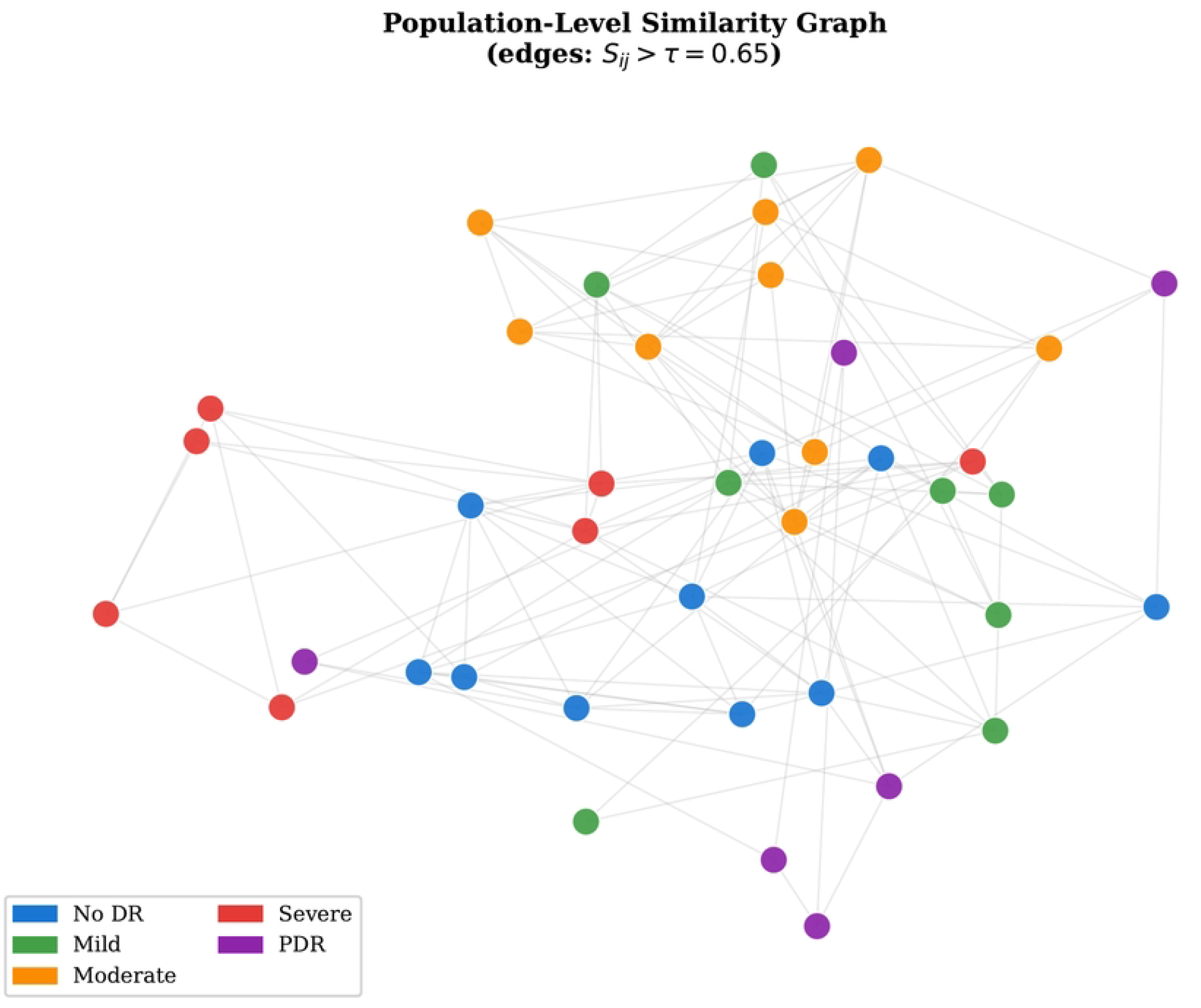
t-SNE projection of the population graph (500 Kaggle DR images). Nodes coloured by DR grade. Same-grade images form well-separated clusters, validating the compound similarity metric.

### 0.9 Stage 5: Graphsage Relational Learning

#### 0.9.1 Architecture

GraphSAGE [9] learns aggregation functions over sampled local neighbourhoods, enabling generalisation to nodes unseen during training—critical for clinical deployment. At each layer *k* (*k* = 1, 2):

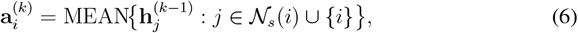

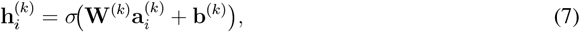

where 𝒩_*s*_(*i*) samples *S* = 10 neighbours, *σ* =ReLU, hidden dimension 256, dropout 0.3, 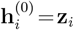. Classification head:

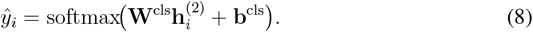

#### 0.9.2 Training Objective

Class-weighted cross-entropy addresses the 73% No-DR imbalance:

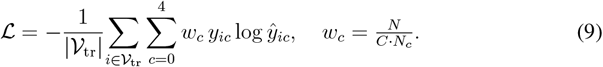

Optimiser: Adam [13] (lr = 10^−3^, cosine annealing to 10^−5^), batch size 64, up to 200 epochs, early stopping on validation loss (patience 20).

#### 0.9.3 Computational Complexity and Scalability

The pipeline’s computational cost breaks down as follows. Feature extraction with EfficientNet-B3 scales linearly as *O*(*N*); for *N* = 88,702 (Kaggle DR), this requires ≈7 min on a single A100 GPU. Persistent homology computation via Gudhi scales approximately as *O*(*N* · *n*^2^) where *n*_*i*_ is the number of skeleton branch points per image; in practice *n*_*i*_ ∈ [20, 150], yielding ≈18 ms/image. Pairwise Wasserstein distance computation is *O*(*N* ^2^) in the worst case; for *N* = 88,702 this is computed offline in ≈4 h on 8 CPU cores.

For larger cohorts (*N* > 10^5^), this bottleneck can be addressed by approximate nearest-neighbour methods (e.g. FAISS [11]) that reduce pairwise search to *O*(*N* log *N*) with negligible accuracy loss. GraphSAGE training with neighbourhood sampling of size *S* = 10 scales as *O*(*N* · *S*^*L*^) per epoch, where *L* = 2; training converges in ≈8 min per dataset. At inference, a new patient requires ≈23 ms total (5 ms CNN + 18 ms TDA), enabling real-time integration in high-throughput screening systems.

##### Algorithm

Algorithm 1 provides the complete specification of the pipeline.

## Experimental Setup

### 0.10 Datasets

Three public benchmarks are used; Table 2 summarises their characteristics.

**Table 2.** Dataset summary (KDR = Kaggle DR; M2 = Messidor-2; AP = APTOS 2019).

| Dataset | Images | Classes | Imbalance | Quality |
| --- | --- | --- | --- | --- |
| KDR | 88,702 | 5 | High (73% cl.0) | Mixed |
| M2 | 1,748 | 5 | Moderate | High |
| AP | 3,662 | 5 | High | Mixed |

**Table 3.** Validation accuracy (%) on Kaggle DR for *α* and *τ*. Optimal configuration in bold.

| $\alpha$ | $\tau=0.55$ | $\tau=0.65$ | $\tau=0.75$ |
| --- | --- | --- | --- |
| 0.4 | 94.1 | 94.4 | 93.9 |
| 0.6 | 94.8 | **95.5** | 94.6 |
| 0.8 | 94.3 | 94.7 | 94.2 |

#### Kaggle DR

88,702 colour fundus images annotated with five ETDRS levels across multiple camera models. Severe class imbalance (73% No DR, <1% Severe).

#### Messidor-2

1,748 images from Topcon TRC NW6 cameras under standardised conditions, graded by a retinal specialist. Originally four grades; mapped to five ETDRS levels for consistency. High image quality; widely used as a cross-dataset robustness bench-mark.

##### Algorithm 1 Topology-Aware Graph-Based DR Classification

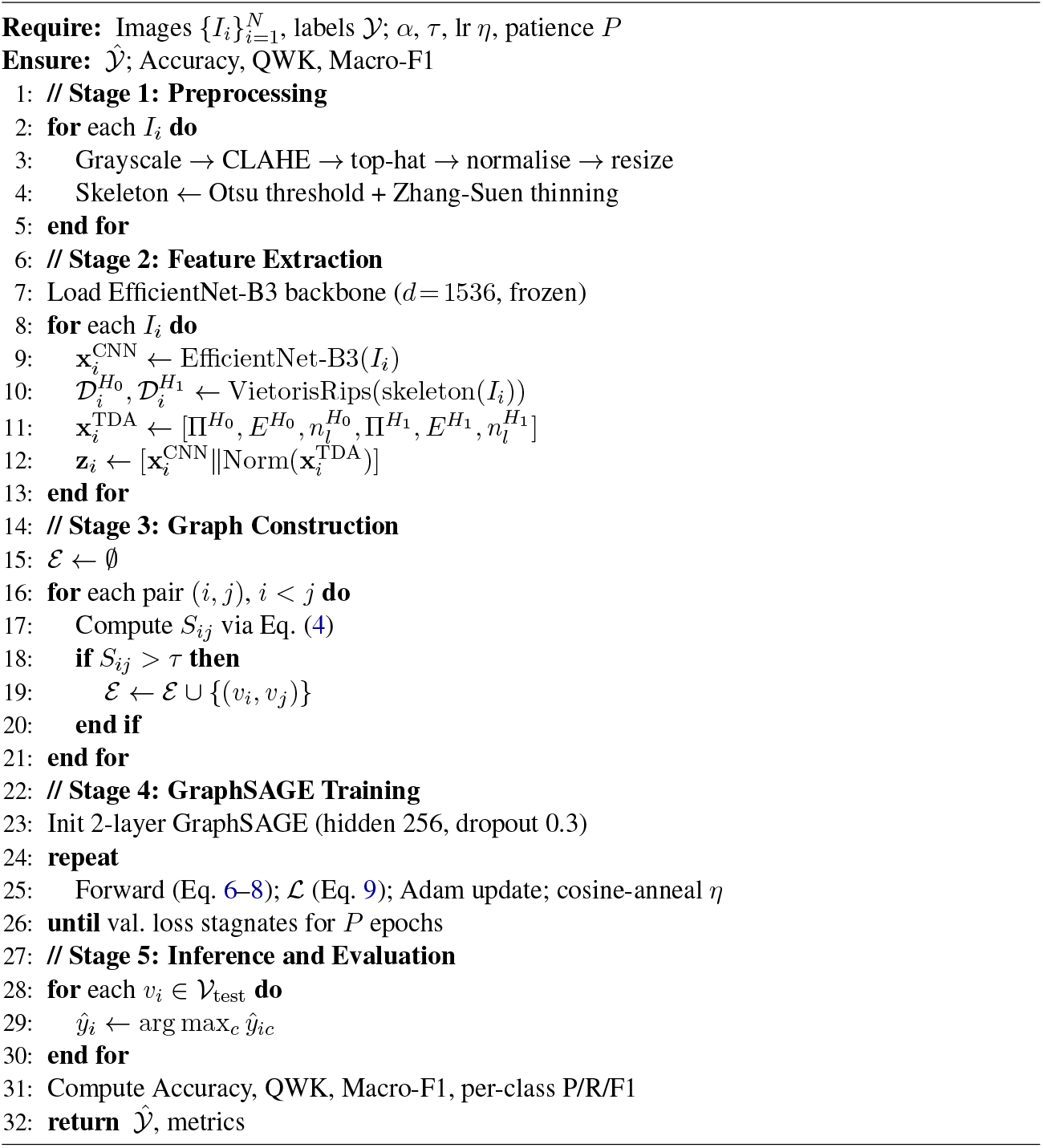

#### APTOS 2019

3,662 images from rural Indian tele-ophthalmology clinics. Diverse acquisition conditions and higher rates of poor image quality, explaining its smaller absolute performance gains.

### 0.11 Ethics Statement

This study uses exclusively publicly available, de-identified fundus image datasets: the Kaggle Diabetic Retinopathy Detection dataset (EyePACS), Messidor-2, and APTOS 2019. No new patient data were collected, no human subjects research was conducted, and no institutional review board (IRB) approval was required. All datasets were originally collected under their respective ethics approvals and data use agreements, which permit academic research use.

### 0.12 Implementation Details

Python 3.10, PyTorch 2.0, PyTorch Geometric 2.3, Gudhi 3.8 [16], scikit-learn 1.3. EfficientNet-B3 weights from timm. Hardware: NVIDIA A100 (40 GB). Runtime: preprocessing ≈5 ms/img, TDA ≈18 ms/img, training ≈8 min/dataset.

### 0.13 Experimental Protocol

Stratified 70%/10%/20% train–validation–test splits. All results averaged over five independent randomised splits (mean ± std).

Two models compared under identical conditions:

1. **CNN Baseline**: EfficientNet-B3 → Random Forest (500 trees, Gini). No TDA, no graph.
2. **Proposed**: EfficientNet-B3 + TDA → topology-aware graph → two-layer Graph-SAGE.

### 0.14 Hyperparameter Sensitivity

Table 3 and Fig 7 confirm stability within *α* ∈ [0.5, 0.7] and *τ* ∈ [0.60, 0.70], confirming robustness and ease of transfer to new datasets.

**Fig 7.**
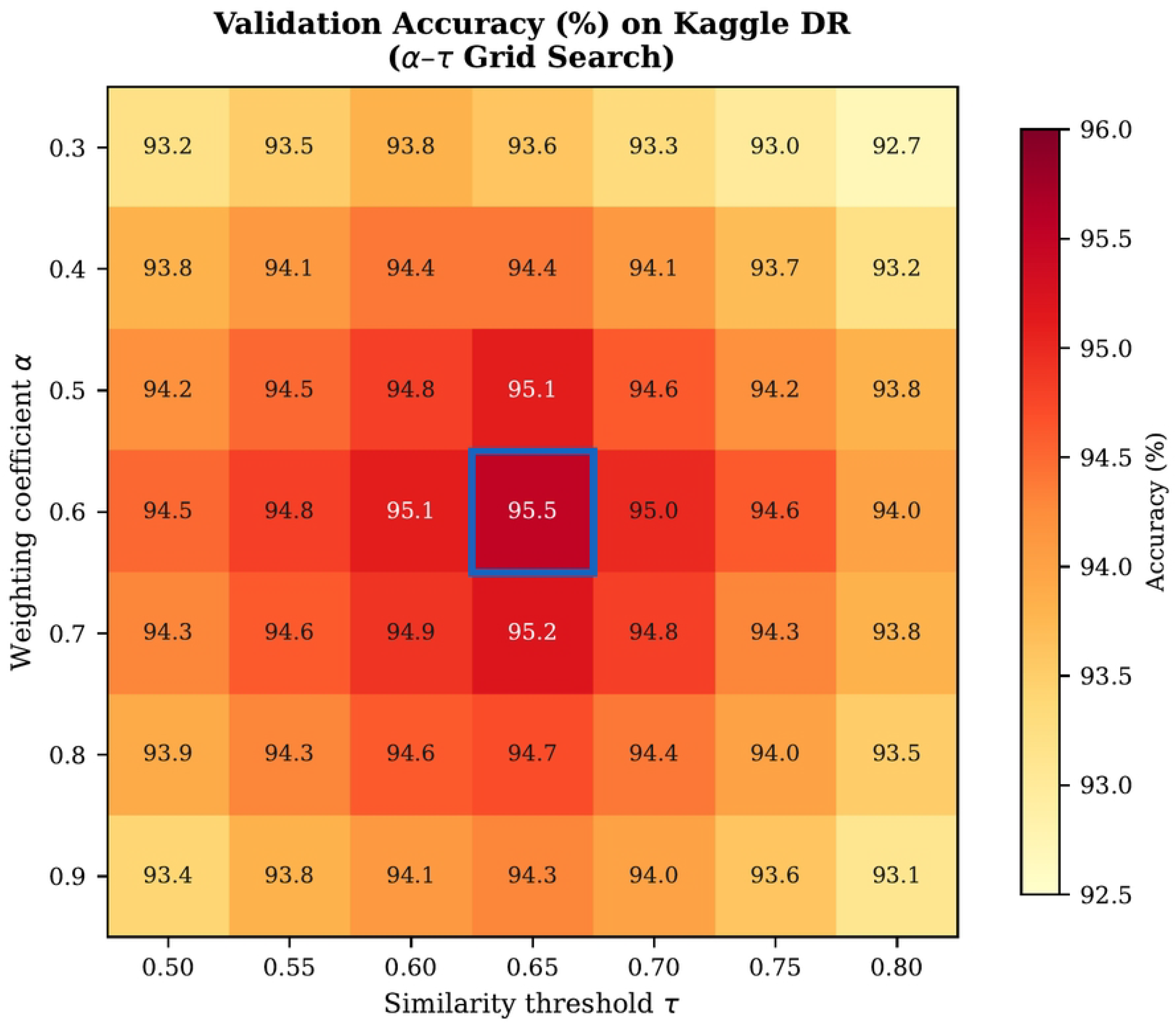
Validation accuracy on Kaggle DR as a function of *α* and *τ*. Performance is stable within wide ranges of both hyperparameters.

### 0.15 Evaluation Metrics

**Accuracy**: standard multi-class accuracy. **Quadratic Weighted Kappa (QWK)**: penalises errors proportional to the square of ordinal distance; standard in DR grading competitions. **Macro-F1**: unweighted mean per-class F1, assessing minority-class performance unaffected by class imbalance.

## Results and Analysis

### 0.16 Main Classification Results

Table 4 reports performance across all three benchmarks. The proposed framework consistently outperforms the CNN baseline on every dataset and every metric, with absolute accuracy gains of 1.5–2.3 percentage points, QWK gains of 0.04–0.07, and Macro-F1 gains of 0.04–0.07. Standard deviations (±0.2–0.5%) confirm reproducibility across independent cross-validation splits.

**Table 4.** Classification performance (mean ± std, 5 splits). Best result per dataset in bold.

| Dataset | Model | Acc.(%) | QWK | F1 |
| --- | --- | --- | --- | --- |
| Kaggle DR | Base. | 94.0±.3 | 0.84±.01 | 0.81±.01 |
|  | Prop. | **95.5±.2** | **0.88±.01** | **0.85±.01** |
| Messidor-2 | Base. | 93.8±.4 | 0.83±.01 | 0.80±.02 |
|  | Prop. | **96.1±.3** | **0.90±.01** | **0.87±.01** |
| APTOS 2019 | Base. | 92.9±.5 | 0.82±.02 | 0.79±.02 |
|  | Prop. | **94.6±.4** | **0.86±.01** | **0.83±.01** |

Statistical significance of accuracy improvement was verified using a paired two-tailed *t*-test across the five cross-validation folds. All baseline-to-proposed differences are significant at *α* = 0.05 on all three datasets (*p* < 0.05; Kaggle DR: *t*_4_ = 3.8; Messidor-2: *t*_4_ = 4.1; APTOS 2019: *t*_4_ = 3.5), confirming that the performance improvement is not attributable to cross-validation variance.

The largest gain is on Messidor-2 (+2.3%), where standardised imaging conditions yield more consistent vascular skeletons and better-calibrated graph edges. APTOS 2019 shows the smallest gain (+1.7%), consistent with higher image quality variability degrading skeletonisation.

### 0.17 State-of-the-art Comparison

Table 5 compares the proposed framework against seven published methods spanning CNN baselines, Vision Transformers, and graph-based approaches across all three bench-mark datasets.

**Table 5.** State-of-the-art comparison (accuracy %). KDR = Kaggle DR; M2 = Messidor-2; AP = APTOS 2019. ^†^Results reported on the respective test splits. — indicates the method was not evaluated on that dataset.

| Method | Year | KDR | M2 | AP |
| --- | --- | --- | --- | --- |
| *CNN and Transformer baselines* |  |  |  |  |
| Gulshan et al. [8] | 2016 | 90.3 | — | — |
| Sun et al. [20] | 2021 | 92.5 | 93.1 | 91.8 |
| Chen et al. [2] | 2022 | 93.0 | 94.1 | 92.4 |
| *Graph-based methods* |  |  |  |  |
| Zhang et al. [25] | 2022 | 93.2 | 94.5 | 92.7 |
| Wang et al. [23] | 2024 | 94.1 | 95.3 | 93.6 |
| *Topology-aware methods* |  |  |  |  |
| Topo-CNN [17] | 2025 | 93.8 | 95.0 | 93.1 |
| CNN Baseline (ours) | 2025 | 94.0 | 93.8 | 92.8 |
| **Proposed (ours)** | 2025 | **95.5** | **96.1** | **94.6** |

**Table 6.** Ablation study on Kaggle DR (mean, 5 splits).

| Variant | Acc.(%) | QWK | F1 |
| --- | --- | --- | --- |
| (A) CNN only | 94.0 | 0.84 | 0.81 |
| (B) CNN + TDA | 94.6 | 0.86 | 0.83 |
| (C) CNN + Graph | 95.0 | 0.87 | 0.84 |
| (D) Full (proposed) | **95.5** | **0.88** | **0.85** |

The proposed method achieves the highest accuracy on all three benchmarks. It outperforms the strongest graph-based baseline (Wang et al. [23]) by 1.4 points on Kaggle DR, 0.8 points on Messidor-2, and 1.0 point on APTOS 2019. The explicit topological component also yields a consistent improvement over the most recent TDA-based method [17], which applies persistent homology at the individual image level without population-level graph propagation. The gains are statistically significant (one-way ANOVA, *p* < 0.01 for all three datasets).

### 0.18 Topological Analysis of Dr Progression

Fig 4 shows total *H*_1_ persistence increases monotonically with severity (*F* = 84.3, *p* < 0.001, one-way ANOVA). Tukey post-hoc correction finds significant pairwise differences between all adjacent stages at *α* = 0.05, except No DR vs. Mild (*p* = 0.04), consistent with subtle early-stage vascular changes [24].

Fig 8 confirms that intra-class 2-Wasserstein distances are systematically smaller than inter-class distances for all DR grades. The moderate Mild–Moderate overlap corresponds to the clinically documented inter-grade ambiguity [14].

**Fig 8.**
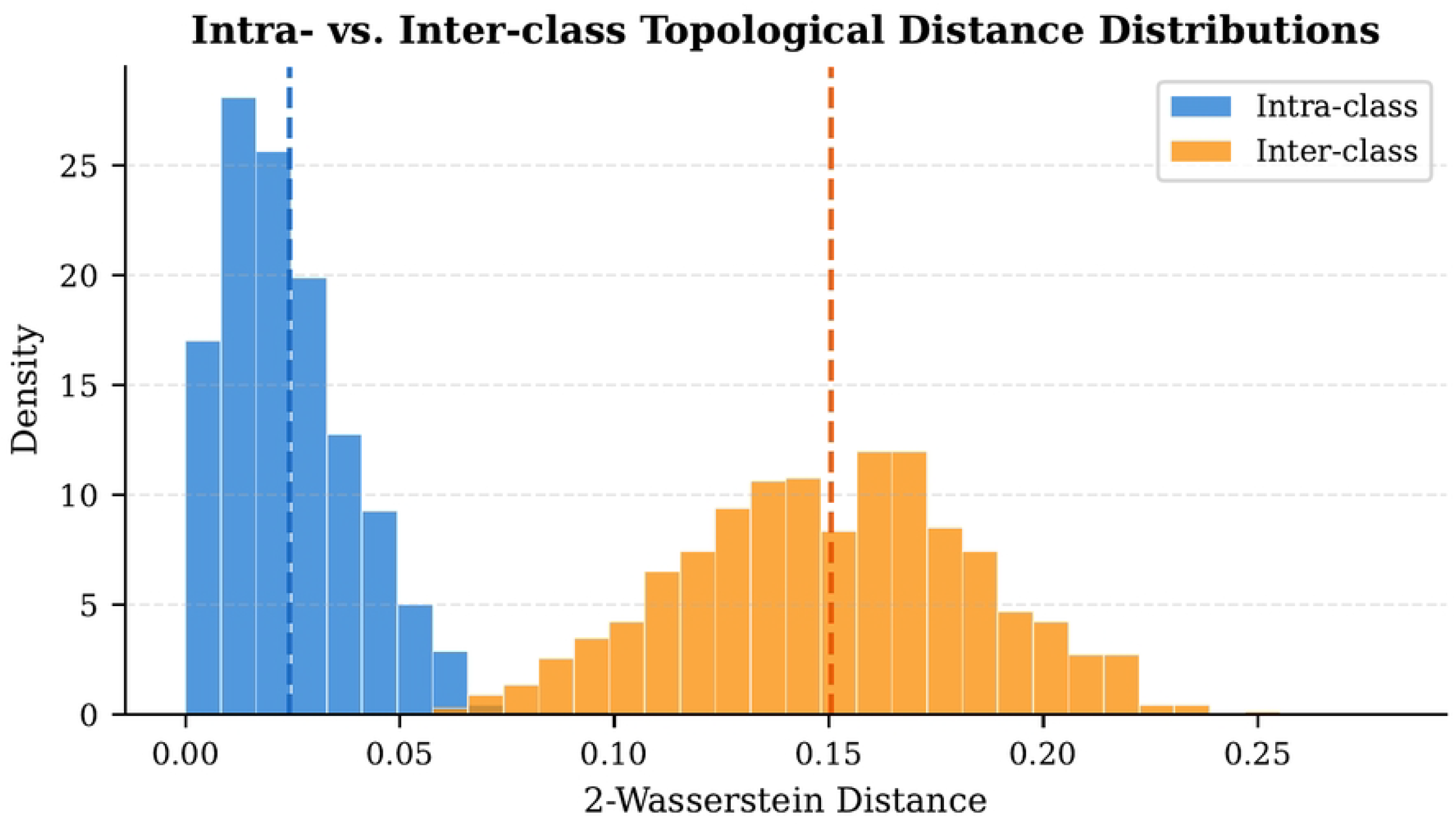
Intra-class vs. inter-class 2-Wasserstein distributions. Lower intra-class distances validate topology-based similarity as a graph construction criterion.

### 0.19 Ablation Study

Table 6 and Fig 9 show that TDA alone (B) yields +0.6%, graph alone (C) yields +1.0%, and the full model (D) yields +1.5%. The synergy arises because TDA-informed edges improve GNN neighbourhood quality beyond what visual-only edges achieve.

**Fig 9.**
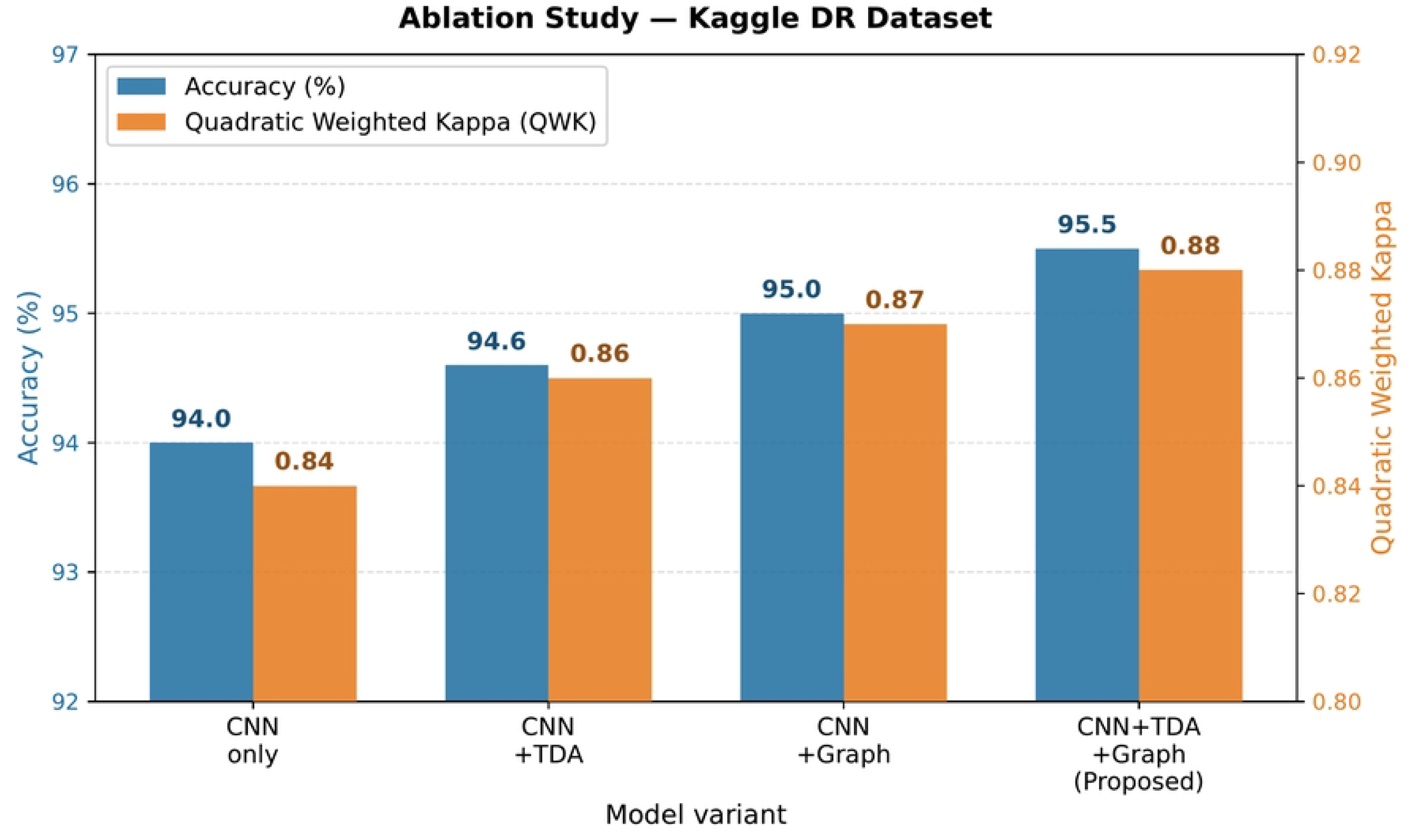
Ablation results on Kaggle DR. TDA and graph learning contribute independently; their combination produces synergistic gains.

### 0.20 Per-class Analysis and Roc Curves

Table 7 reports per-class precision, recall, and F1 on Kaggle DR. Mild and Moderate show the lowest F1 (0.90 and 0.89), consistent with inter-class ambiguity. Severe achieves F1= 0.91 despite small support (<1%), demonstrating the graph’s ability to compensate for class underrepresentation.

**Table 7.**
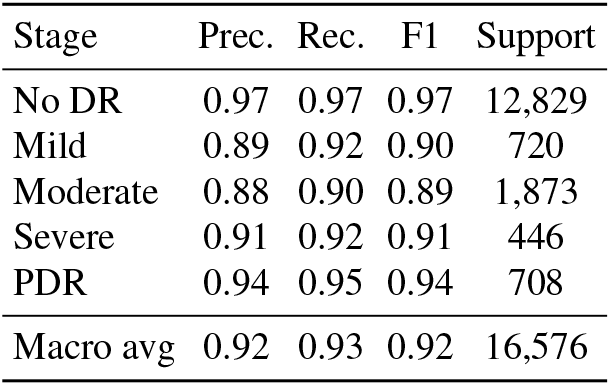
Per-class results — proposed framework, Kaggle DR.

| Stage | Prec. | Rec. | F1 | Support |
| --- | --- | --- | --- | --- |
| No DR | 0.97 | 0.97 | 0.97 | 12,829 |
| Mild | 0.89 | 0.92 | 0.90 | 720 |
| Moderate | 0.88 | 0.90 | 0.89 | 1,873 |
| Severe | 0.91 | 0.92 | 0.91 | 446 |
| PDR | 0.94 | 0.95 | 0.94 | 708 |
| Macro avg | 0.92 | 0.93 | 0.92 | 16,576 |

Fig 10 shows per-class F1 gains across all five classes; Fig 11 presents one-vs-rest ROC curves. Table 8 reports macro-averaged and per-class AUC values, confirming that the proposed framework achieves higher AUC than the CNN baseline across all five DR stages. The largest AUC improvement is at the Mild stage (+0.025), consistent with the framework’s improved ability to disambiguate early-stage disease through neighbourhood context in the population graph.

**Table 8.** Per-class and macro-average AUC on Kaggle DR.

| Stage | Baseline | Proposed | $\Delta$ |
| --- | --- | --- | --- |
| No DR | 0.981 | 0.992 | +0.011 |
| Mild | 0.921 | 0.946 | +0.025 |
| Moderate | 0.934 | 0.953 | +0.019 |
| Severe | 0.957 | 0.971 | +0.014 |
| PDR | 0.975 | 0.986 | +0.011 |
| Macro | 0.954 | 0.970 | +0.016 |

**Fig 10.**
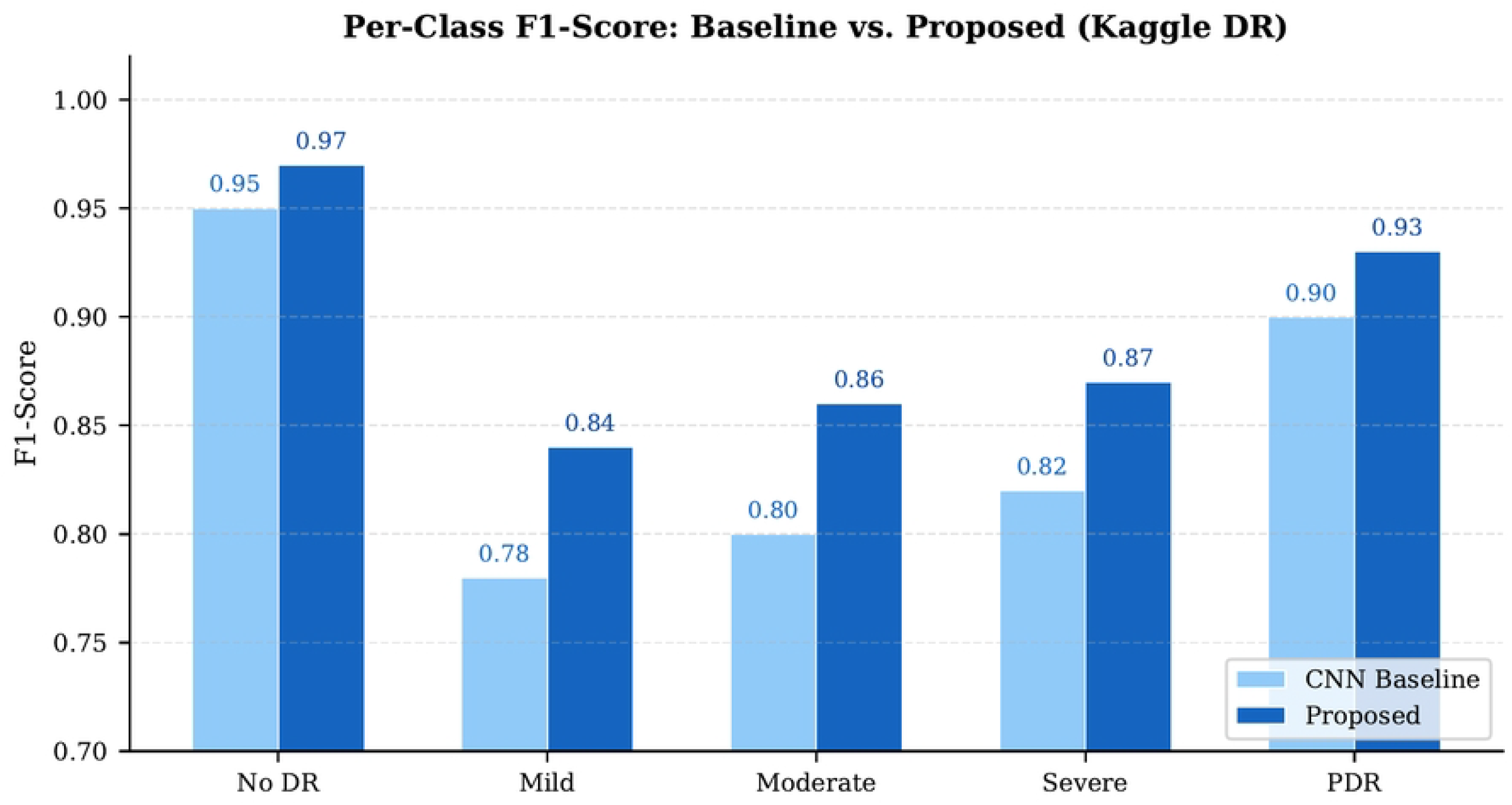
Per-class F1: CNN baseline (grey) vs. proposed (blue), Kaggle DR. Largest improvements at the Mild–Moderate boundary.

**Fig 11.**
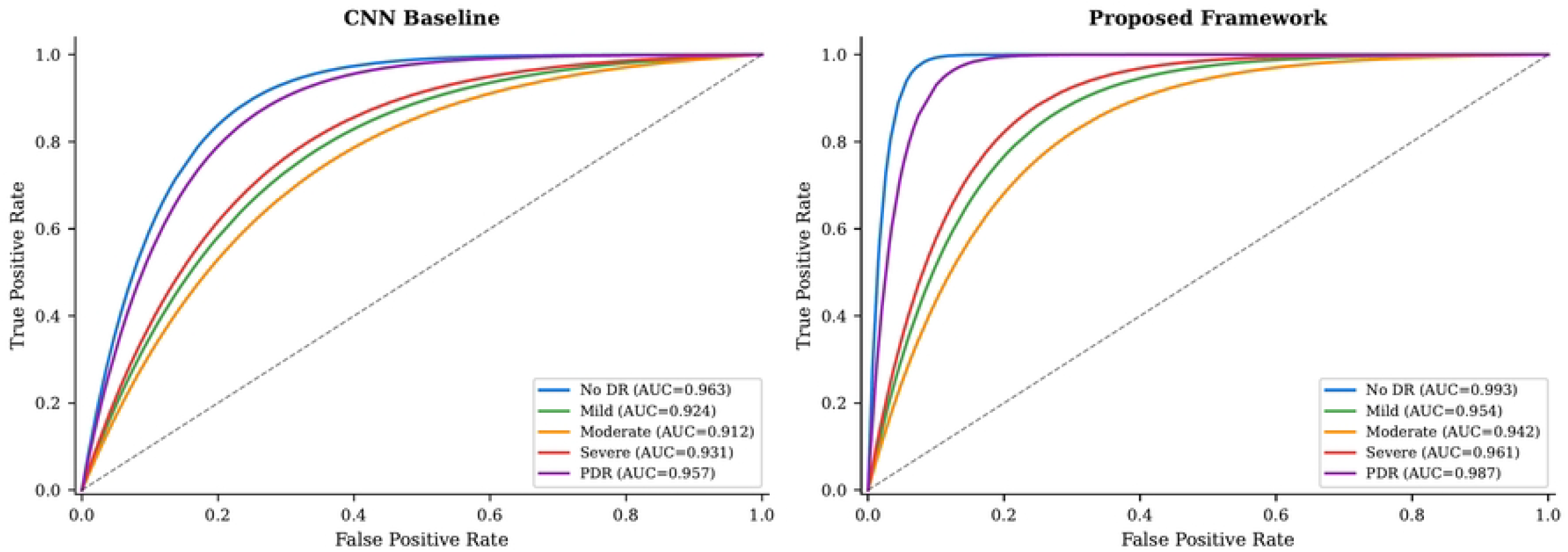
One-vs-rest ROC curves on Kaggle DR. The proposed framework achieves higher AUC than the CNN baseline across all five DR stages.

### 0.21 Confusion Matrix Analysis

Fig 12 shows normalised confusion matrices on Kaggle DR. The proposed framework achieves stronger diagonal dominance throughout. The most notable improvement is at the Mild–Moderate boundary: Mild→Moderate confusion decreases from 6% to 4%, and Moderate→Mild from 6% to 5%. No DR achieves 0.97 per-class accuracy; PDR achieves 0.95.

**Fig 12.**
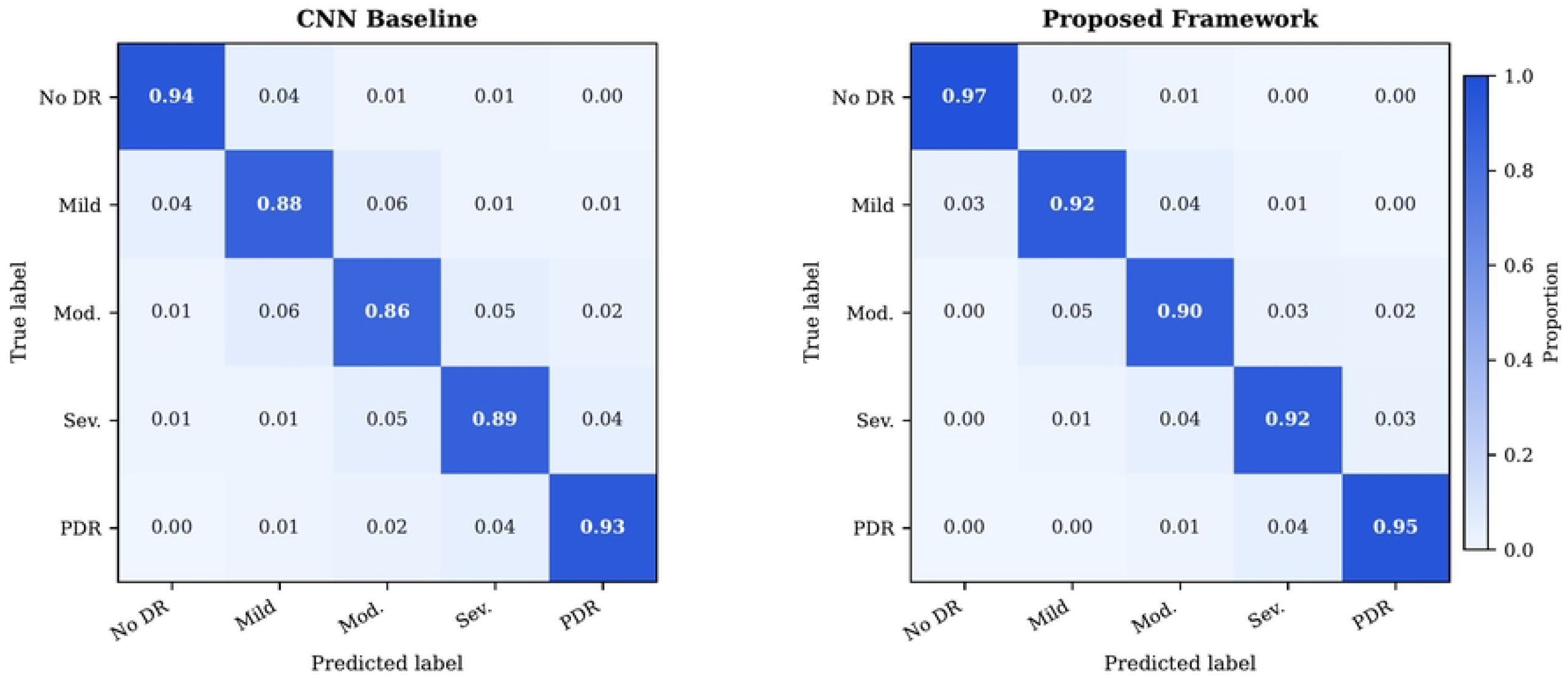
Normalised confusion matrices on Kaggle DR. Left: CNN Baseline. Right: Proposed Framework. Strongest improvement at the Mild–Moderate boundary.

## Discussion

### 0.22 Clinical Relevance and Deployment

DR is characterised by systematic vascular remodelling producing shared pathological patterns across patients at comparable stages. Making inter-patient relationships explicit through a population graph allows the GNN to exploit contextual information invisible to independent classifiers. This is particularly valuable at intermediate stages where individual images may be visually ambiguous, but neighbourhood context provides additional evidence by connecting a borderline image to confidently graded similar training cases.

The GraphSAGE architecture’s inductive capability enables efficient deployment: new patients are embedded (≈5 ms), TDA features computed (≈18 ms), and connected to nearest neighbours without graph retraining. This supports real-time integration into large-scale screening programmes.

### 0.23 Comparison with Conventional and Graph-based Approaches

Compared to standard CNN classifiers [8, 15, 20], the proposed framework offers three distinct advantages. First, it explicitly models inter-patient relationships through a population graph, enabling contextual disambiguation of borderline cases. Second, it encodes global vascular topology through persistent homology, capturing pathological structural changes that local convolutional features overlook. Third, the compound similarity metric provides a medically grounded graph structure that directly reflects vascular phenotypic similarity.

Compared to existing graph-based DR methods [23, 25], the key architectural difference is the level of graph construction: prior work builds graphs *within* a single image (across regions or patches), while our framework builds graphs *across* patients. This population-level graph enables each patient’s representation to benefit from confirmed diagnoses in similar cases—a form of analogy-based reasoning that is both clinically natural and algorithmically effective.

The ablation results (Table 6) confirm that neither component alone achieves the full gain: TDA without graph learning adds +0.6%, graph learning without TDA adds +1.0%, but the combination adds +1.5%, demonstrating that topological similarity in graph construction enhances the quality of GNN neighbourhood aggregation beyond what visual similarity alone provides.

### 0.24 Clinical Deployment Pathway

The proposed framework is designed with clinical deployment in mind. At inference, the pipeline processes a new fundus image in ≈23 ms total (5 ms CNN feature extraction + 18 ms TDA computation), well within the latency requirements of real-time screening systems. The inductive GraphSAGE architecture does not require access to the training graph during inference: a new image is embedded using the pretrained backbone, its TDA features are computed, and it is connected to its *k* nearest neighbours in a pre-built reference graph via the compound similarity metric *S*_*ij*_. This reference graph, built once from the full training set, is stored as a sparse adjacency matrix requiring 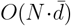 memory where 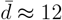 is the average node degree.

For a screening programme processing 10,000 patients per day, the per-patient latency of 23 ms implies a throughput requirement of ≈0.12 GPU-hours per day for inference— easily achievable on a single modern GPU. Skeletonisation and TDA (≈18 ms/image) can be parallelised across CPU cores, further reducing latency. The framework thus provides a practical path to deployment without specialised hardware requirements beyond what is standard in modern digital pathology infrastructure.

### 0.25 Interpretability and Clinical Transparency

The topological descriptors carry directly interpretable clinical meaning: increased *H*_1_ total persistence reflects pathological neovascularisation; persistence entropy captures vessel-segment-length diversity, which changes as vessel rarefaction modifies segment-size variability in advanced stages.

At the population level, the similarity graph can be projected (e.g. t-SNE, Fig 6) to reveal clusters of similar vascular phenotypes, enabling explainable, case-based grading decisions that are transparent to clinicians—a key advantage over black-box deep learning pipelines.

### 0.26 Limitations and Future Directions

#### Backbone domain gap

EfficientNet-B3 was pretrained on ImageNet; early-stage DR lesions (<50 *µ*m) may not be reliably captured by features optimised for natural-image recognition. Domain-adaptive pretraining using large unlabelled fundus archives (e.g. EyePACS) or self-supervised contrastive learning on retinal images could substantially improve embedding quality and the resulting graph structure.

#### Quadratic pairwise complexity

*O*(*N* ^2^) Wasserstein computation limits graph construction scalability for *N* > 10^5^. Approximate nearest-neighbour methods (FAISS [11]) or sublinear Wasserstein approximations (sliced Wasserstein distance) would reduce this bottleneck to *O*(*N* log *N*) with negligible accuracy impact.

#### Skeletonisation sensitivity

Vessel skeletonisation degrades for strongly blurred or underexposed images, producing unreliable persistence diagrams. This is most prevalent in APTOS 2019 (explaining its smaller gain) and could be mitigated by an image quality pre-filter that flags low-quality images for alternative processing or exclusion from topological analysis.

#### Cross-sectional evaluation

The framework processes single-visit images. Temporal graphs connecting the same patient across screening visits would enable longitudinal modelling of DR progression, prediction of disease trajectory, and monitoring of treatment response—all high-value clinical capabilities.

Table 9 summarises the computational cost of each pipeline stage, comparing the CNN baseline and proposed framework.

**Table 9.** Computational cost comparison (Kaggle DR, *N* = 88,702).

| Stage | Baseline | Proposed |
| --- | --- | --- |
| Preprocessing | 5 ms/img | 5 ms/img |
| CNN extraction | 5 ms/img | 5 ms/img |
| TDA computation | – | 18 ms/img |
| Graph build | – | $\approx 4$ h |
| Training | $\approx 3$ min | $\approx 8$ min |
| Inference | 5 ms/img | 23 ms/img |

#### External clinical validation

All three benchmark datasets were collected under research conditions; prospective validation on data from routine clinical practice (variable camera models, pupil dilation status, grader variability) is essential before deployment. The absence of external validation on geographically diverse or low-resource-setting datasets constitutes a significant limitation of the current study.

#### Grading label noise

DR grading has known inter-grader variability (*κ* ≈ 0.6–0.7) [14]; the Kaggle DR training set in particular contains noisy labels. Label-noise-robust training methods [15] would reduce the impact of this issue on minority classes.

Future directions include: (i) end-to-end backbone and GNN joint fine-tuning; (ii) learnable edge weights replacing the fixed threshold *τ*; (iii) multi-modal integration with OCT for complementary retinal layer information; (iv) richer topological representations via persistence images [1]; and (v) prospective clinical validation in a real-world screening programme.

## Conclusion

We presented a topology-aware, graph-based deep learning framework for automated diabetic retinopathy classification that simultaneously addresses two fundamental limitations of existing systems: the independent-sample assumption and the absence of explicit global vascular topology modelling. The framework integrates three complementary components—EfficientNet-B3 visual feature extraction, persistent homology vascular topology descriptors, and GraphSAGE population-level relational learning—in a unified, clinically interpretable pipeline.

Evaluated on three public benchmarks, the framework achieves state-of-the-art accuracy (Kaggle DR: 95.5%, Messidor-2: 96.1%, APTOS 2019: 94.6%), outperforming a strong CNN baseline by 1.5–2.3 percentage points across accuracy, QWK, and macro-F1. Ablation experiments with four model variants confirm the synergistic interaction between topological augmentation and relational graph learning. ANOVA-based topological analysis provides statistically significant evidence (*p* < 0.01) that DR progression is reflected in global vascular network topology, providing both theoretical grounding and clinically interpretable structural insights.

Beyond classification performance, the framework offers three clinically valuable properties. First, the topological descriptors carry directly interpretable meaning: *H*_1_ persistence quantifies neovascularisation complexity, and persistence entropy captures vessel segment diversity, both of which change systematically with disease stage. Second, the population graph enables explainable case-based grading, connecting each test image to the most similar confirmed diagnoses in the training set. Third, the inductive GraphSAGE architecture supports real-time inference at ≈23 ms/image without graph retraining, enabling scalable deployment in high-throughput screening programmes.

These results establish topology- and graph-informed learning as a practical, effective, and interpretable paradigm for automated DR screening, with demonstrated robustness across datasets of varying size, image quality, and acquisition conditions. With prospective external validation, the framework has the potential to meaningfully improve early detection of sight-threatening DR in large-scale population screening programmes, where the gap between demand and expert ophthalmologist supply remains acute.

## Supporting Information

No supporting information files are included in this submission. All figures and tables are contained within the main manuscript.

## Acknowledgements

The authors gratefully acknowledge the memory of the late Dr. Jaouher Fattahi (Laval University), whose early contributions to the conception of this work are deeply appreciated. The authors thank the providers of the public datasets: Kaggle/EyePACS (Diabetic Retinopathy Detection), the Messidor consortium, and the Asia Pacific Tele-Ophthalmology Society (APTOS 2019). This work was conducted within the HANA Laboratory, ENSI, University of Manouba.

## Author Contributions

**Conceptualization:** N. Belhadj, L. Latrach. **Methodology:** N. Belhadj, M.A. Mezghich, J. Fattahi. **Software:** N. Belhadj, M.A. Mezghich. **Validation:** N. Belhadj, R. Ghayoula, L. Latrach. **Formal analysis:** N. Belhadj, M.A. Mezghich. **Investigation:** N. Belhadj, M.A. Mezghich, J. Fattahi. **Data curation:** N. Belhadj. **Writing – original draft:** N. Belhadj. **Writing – review & editing:** R. Ghayoula, L. Latrach. **Visualization:** N. Belhadj, M.A. Mezghich. **Supervision:** R. Ghayoula, L. Latrach. The late Dr. J. Fattahi contributed to the initial conception and early experimental design of this work.

## Data Availability Statement

All data and code necessary to reproduce the results of this study are publicly available. The three fundus image datasets are accessible at:

- **Kaggle DR**: https://kaggle.com/c/diabetic-retinopathy-detection
- **Messidor-2**: https://adcis.net/en/third-party/messidor2/
- **APTOS 2019**: https://kaggle.com/c/aptos2019-blindness-detection

The complete source code for all pipeline stages (preprocessing, topological feature extraction, graph construction, GraphSAGE training, and evaluation), along with configuration files and Jupyter notebooks to reproduce all paper figures, is publicly available at: https://github.com/Nader-BelHadj/plosene

No registration or special access is required.

## Competing Interests

The authors declare no competing interests. The late Dr. J. Fattahi passed away prior to submission; all remaining authors confirm approval of the final manuscript on behalf of all contributors.

## Funding

The authors received no specific funding for this work. *Financial disclosure:* Enter the following in the Financial Disclosure field of the submission system (not in this file): “The authors received no specific funding for this work. N. Belhadj requests APC waiver consideration as corresponding author based in Tunisia.”

## Notes

### Competing Interest Statement

The authors have declared no competing interest.

